# Maximizing the productivity of co-transcriptional in vitro transcription

**DOI:** 10.64898/2026.07.28.740954

**Authors:** Wei He, Kunkun Zhang, Guiying Ji, Hongli Zhou, Ya Dai, Minghao Xu, Xiaoyu Xu, Qiuheng Jin

**Affiliations:** Vazyme Biotech Co., Ltd. Nanjing 210038, China; College of Biology and the Environment, Nanjing Forestry University, Nanjing 210037, China; BioGeometry, Beijing 100083, China; Academy for Advanced Interdisciplinary Studies, Peking University, Beijing 100871, China

## Abstract

m*RNA* therapeutics have demonstrated transformative potential in infectious disease, oncology and genetic disorders. Parallel to the rapid clinical advancement of mRNA therapies, research into in vitro transcription (IVT), the pivotal process of mRNA manufacturing, has intensified significantly. However, the majority of these studies omitted cap analog, an essential substrate for co-transcriptional capped IVT (co-IVT), likely to simplify experimental systems. Nevertheless, co-IVT remains the dominant industrial standard for the production of mRNA therapeutics. Here, we report an optimized co-IVT process that achieves a yield of more than 26 g/L while maintaining capping efficiency of over 99.5%, and achieving a 30-fold reduction in double-stranded RNA (dsRNA), the most critical process-related impurity. These performance gains substantially improve mRNA production efficiency and enhance mRNA quality, thereby accelerating the transition to commercial-scale manufacturing of mRNA therapeutics. Moreover, the performance was attained using either of two mechanistically distinct engineered T7 RNA polymerase (RNAP) mutants. Both mutants enabled robust mRNA synthesis under process conditions that have not been previously reported. These observations provide mechanistic insights into IVT and advance our fundamental understanding of this critical bio-manufacturing step.

## INTRODUCTION

For enzyme-catalyzed processes to achieve economic viability on an industrial scale, high substrate loading is essential^[1-3]^ . Elevated substrate concentrations enhance volumetric productivity and final product titers, thereby lowering downstream processing costs and overall manufacturing expenditures. However, the optimization of enzymatic reaction conditions is constrained by the solubility limits of the substrate or product. In some cases, process developers employ specialized solvents to extend these solubility boundaries, enabling higher substrate loading or improved product accumulation^[4, 5]^ .

Most naturally occurring enzymes have evolved under physiological conditions and, therefore, exhibit substrate inhibition at industrially relevant high-substrate concentrations. This limitation markedly suppresses reaction rates and hinders process intensification^[6-8]^ . Beyond substrate inhibition, high product accumulation imposes a second critical constraint on biocatalytic efficiency. Specifically, product molecules can either inhibit an enzyme by binding to its active or allosteric sites or induce unfavorable shifts in local pH, ionic strength, or solvent polarity. These effects ultimately result in incomplete substrate conversion^[9-11]^, which not only wastes feedstock but also reduces volumetric and molar product yields.

mRNA is a clinically effective therapeutic platform. Prophylactic vaccines against COVID-19 and RSV have received regulatory approval for clinical application, and an influenza candidate is currently under BLA review^[12]^ . Concurrently, a range of emerging therapeutic modalities, including individualized cancer immunotherapy^[13, 14]^, in vivo cell reprogramming^[15]^, and protein replacement therapy^[16]^, have achieved clinical proof-of-concept. A recent industry landscape analysis reported that 244 therapeutic Mrna candidates are in clinical development, excluding those targeting COVID-19^[17]^ . The demand for mRNA drugs, driven by these clinical pipelines and preclinical research activities currently generates hundreds of millions of dollars in annual revenue and is projected to grow rapidly. Despite this accelerating demand, the mRNA manufacturing process remains significantly under-optimized.

mRNA is produced via in vitro transcription (IVT), a process catalyzed by bacteriophage RNA polymerases (RNAPs), predominantly T7 RNAP. In the presence of Mg^2+^ as a cofactor, T7 RNAP specifically binds to its cognate promoter sequence on a linearized DNA template and initiates de novo RNA synthesis, producing a transcript complementary to the DNA antisense strand^[18]^ . Virtually all clinically relevant mRNAs incorporate a 5′-cap structure, either added post-transcriptionally through a multi-enzymatic reaction or synthesized co-transcriptionally during IVT by introducing a fifth substrate, a cap analog that serves as a primer^[19, 20]^ . The latter approach, commonly termed co-transcriptional IVT (co-IVT), has emerged as the predominant strategy owing to its superior overall process efficiency.

IVT is widely recognized for its high efficiency, enabling the rapid synthesis of 6–10 g/L mRNA within several hours. However, the yield is far below the solubility limits of the substrate and product. NTPs exhibit aqueous solubilities of at least 200 mM aqueous solubility, resulting in a theoretical RNA yield exceeding 65 g/L. The solubility of mRNA is highly dependent on sequence composition and solvent conditions. Notably, even the least soluble type of RNA (47×CAG) achieved a soluble concentration of 16.3 g/L in Mg^2+^ - containing buffer and remained unprecipitated^[21]^ . Therefore, IVT exhibits substantial untapped potential for yield improvement.

Cost-of-goods (CoGs) analysis identified IVT as the costliest unit operation in mRNA manufacturing, with RNAPs, cap analogs, and DNA templates representing the primary cost drivers^[22]^ . Despite this economic bottleneck, systematic efforts to improve IVT yields remain scarce. Although fed-batch strategies have been explored to enhance the utilization efficiency of RNAP and DNA templates^[23, 24]^, they concurrently lead to disproportionate consumption of cap analogs, thereby offsetting potential cost savings and yielding only marginal improvements in overall process economics. The highest reported IVT yield under batch conditions was 24.9 g/L^[25]^ . However, this process omitted co-transcriptional capping using a cap analog, thereby introducing an additional enzymatic capping unit operation that reduces overall mRNA production efficiency and increases the CoGs.

In this study, we employed two distinct engineered T7 RNAPs to establish a robust, highly productive co-IVT batch process. Critically, this process demonstrated industrial scalability, met the growing demand for therapeutic mRNA, and reduced cap analog consumption without compromising capping efficiency, thereby substantially improving the cost-effectiveness of the IVT reaction.

## MATERIAL AND METHODS

### Reagents

Commercial kits (including high-yield RNA transcription, dsRNA quantification, site-directed mutagenesis, and BCA protein assay kits), enzymes (such as RNase inhibitor [RI], inorganic pyrophosphatase [PPase], and DNase I), and nucleotides (ATP/GTP/CTP/UTP at 100 mM) were supplied by Vazyme Biotech Co., Ltd. (Nanjing, China). Nucleotides (200 mM) were obtained from Wuhu Huarun Biotechnology (Wuhu, China). Mg(Ac)_2_, dithiothreitol, and other chemical reagents were purchased from Aladdin Industrial Corporation (Shanghai, China).

### Generation of T7 RNAP variants

In this study, we leveraged the GeoFlow V2 generative protein design model of BioGeometry^[26]^ to rationally engineer T7 RNAP variants with attenuated Mg^2+^ requirements and improved catalytic performance. GeoFlow V2 excels at integrating atomic-level protein structural data, evolutionary sequence constraints, and physics-informed energy functions to identify beneficial mutations that modulate enzyme–substrate interactions and cofactor-binding affinity. We used this platform to conduct a systematic in silico mutagenesis scan across the catalytic core and Mg^2+^-binding regions of T7 RNAP, generating a curated list of high-potential single-, double-, and triple-point mutations for subsequent experimental screening.

### In vitro mRNA synthesis

The T7 RNAP WT or variants were added to the reaction mixture at a final concentration of 20 U/μL, adjusted according to their specific enzymatic activity. The IVT reaction system contained the following final concentrations: 40 mM HEPES (pH 7.0), 2.5 mM TCEP, 2 mM spermidine, 50 ng/μL DNA template (Fig S1), 4 U/μL RI, 5 × 10^−3^ U/μL PPase, 5–20 mM NTPs, and 5–90 mM magnesium acetate. The capping analogs were included for a co-transcriptional capping at a final concentration of 5–20 mM. The reactions were incubated at 37°C for 2–4 h (Table S1). Upon completion, the resulting mRNA products were collected for downstream purification.

### Small-scale mRNA purification

Following IVT, the reaction mixture was diluted 10-fold with ddH_2_O and incubated at 37°C for 5 min. VAHTS RNA Clean Beads (Vazyme) were then added at a volume ratio of 1.8× relative to that of the diluted sample. The sample was thoroughly mixed via pipetting, incubated for 5 min, and then placed on a magnetic stand for 5 min. The supernatant was discarded, and the beads washed twice with 200 μL of 80% ethanol. After air-drying the beads for 10 min, the tube was removed from the magnetic stand, and 50 μL ddH_2_O was added. The sample was resuspended via pipetting, incubated for 5 min, and placed back onto the magnetic stand for 5 min, whereafter the supernatant containing the purified mRNA was collected. The final mRNA concentration was determined using a NanoDrop spectrophotometer.

### Scaled-up mRNA purification

After the completion of IVT, EDTA-2Na was added at a final concentration of 0–100 mM to ensure complete dissolution. The solution was transferred to a large-volume vessel and diluted to 40 mL with loading buffer (20 mM Tris, 500 mM NaCl, 1 mM EDTA, pH 7.5). The mixture was incubated at 37°C with shaking at 220 rpm for 10 min. For the third and fourth experimental groups, the IVT mixtures were separately diluted to 20 and 40 mL with loading buffer, respectively, and subjected to the same incubation conditions. Upon completion of sample preparation, all samples were loaded onto a 15 mL POROS Oligo (dT)25 affinity column (Thermo Fisher Scientific, Waltham, MA, USA). After the unbound IVT components had been fully eluted through the column and the UV260 signal stabilized, the column was washed with loading buffer. Purified mRNA was subsequently eluted with ddH_2_O. The concentration of eluted mRNA was quantified using a NanoDrop spectrophotometer.

### Capping efficiency analysis

A total of 1.25 μM mRNA product was mixed with the detection probe at a 1:1 ratio and annealed. Following annealing, 25 pmol mRNA per reaction was digested with 20 U RNase H at 25°C for 20 min. For every 25 pmol of mRNA used, 9 mL streptavidin-coated magnetic beads were added. Initially, the beads were placed on a magnetic stand for 5 min until the solution clarified, after which the supernatant was discarded and the beads washed twice with 200 μL ddH_2_O. The RNase H-digested samples were mixed with the beads via pipetting and then incubated for 30 min with gentle rotation. The tube was placed on a magnetic stand for 5 min, the supernatant discarded, and the beads washed twice with wash buffer (5 mM Tris-HCl, 0.5 mM EDTA, 1 M NaCl, pH 7.5), followed by washing twice with ddH_2_O. Finally, the appropriate volume of elution buffer (1% methanol in ddH_2_O) was added to achieve a final mRNA concentration of 1.25 μM. The mixture was vortexed, incubated at 85°C for 3 min, and immediately placed on a magnetic stand. The supernatant was collected for mass spectrometry (MS) analysis.

The MS parameters were as follows: mobile phase A, 0.065% HFIP + 0.0375% DIPEA in water; mobile phase B, 0.065% HFIP + 0.0375% DIPEA in methanol; gradient, 0–20 min from 5% □ 20% B; column, Nano ChromCore C18 (50 × 4.6 mm, 1.8 μm); injection volume, 5 μL; flow rate, 0.3 mL/min; column temperature, 60°C; m/z range, 500–3000; instrument, Thermo Vanquish Flex UPLC coupled to an Orbitrap Exploris 120 MS (Thermo Fisher Scientific).

### mRNA integrity and dsRNA quantification

A total of 100–200 ng mRNA product was diluted to 5–10 ng/μL using the provided dilution buffer (RNA Cartridge Kit #R1C405110; BiOptic Inc., Flintridge, CA, USA). The samples were denatured by incubation at 70°C for 5 min, followed by immediate cooling on ice at 0°C for 5 min. The mRNA integrity was assessed using capillary electrophoresis.

The dsRNA content was quantified using a dsRNA Quantification Kit (Vazyme Biotech Co., Ltd.). This kit employs a double-antibody sandwich ELISA method for the quantitative detection of dsRNA in the IVT system.

### Design of the DoEs

DoEs were created and analyzed using JMP PRO software. Responses included the mRNA yield (mg/mL), capping rate (%), and dsRNA content (ng/mg). With regards to the response to yield and capping rate, four main effects were ranked based on their estimated relative impact on the process output among the ten most influential model terms (Fig S2). The NTP concentration (20 mM each) and reaction time (4 h) were set as non-variable factors in the assay. The response surfaces, effect summary, and prediction profiler data were calculated using built-in functions of the JMP PRO software.

## RESULTS

### Optimization of co-IVT

A recent study reported an IVT yield of 24.9 g/L^[25]^. To evaluate whether this high-yield protocol can be directly adapted to the co-IVT reaction, we reproduced the core process conditions while systematically assessing the impact of the cap analog. Specifically, we replaced the original DNA template with one suitable for co-IVT; supplemented the reactions with three distinct concentrations of the trinucleotide cap analog, m^7^GpppA_m_pG (hereafter designated as the GAG cap); and quantified the mRNA yield and capping rate. We failed to achieve the previously reported yield, even at the lowest tested GAG cap concentration (5 mM). As expected, the mRNA yields progressively decreased with increasing GAG cap concentrations, whereas the capping rate only increased marginally. Critically, even at the highest GAG cap concentration (20 mM), the capping rate remained below the 95% minimum threshold specified for therapeutic mRNA (Fig 1A, B). Subsequently, we applied a design of experiments (DoE) approach to define a feasible design space for the co-IVT reaction that simultaneously would meet the target yield (25 g/L) and capping rate specification (>95%). In both the yield and capping rate models, Mg^2+^ and GAG cap concentrations emerged as the two most statistically significant factors, exerting opposing effects on the two quality attributes (Fig S2). Consequently, no region within the explored parameter space simultaneously achieved both targets, revealing an intrinsic process trade-off (Fig 1C, D).

**Figure 1:**
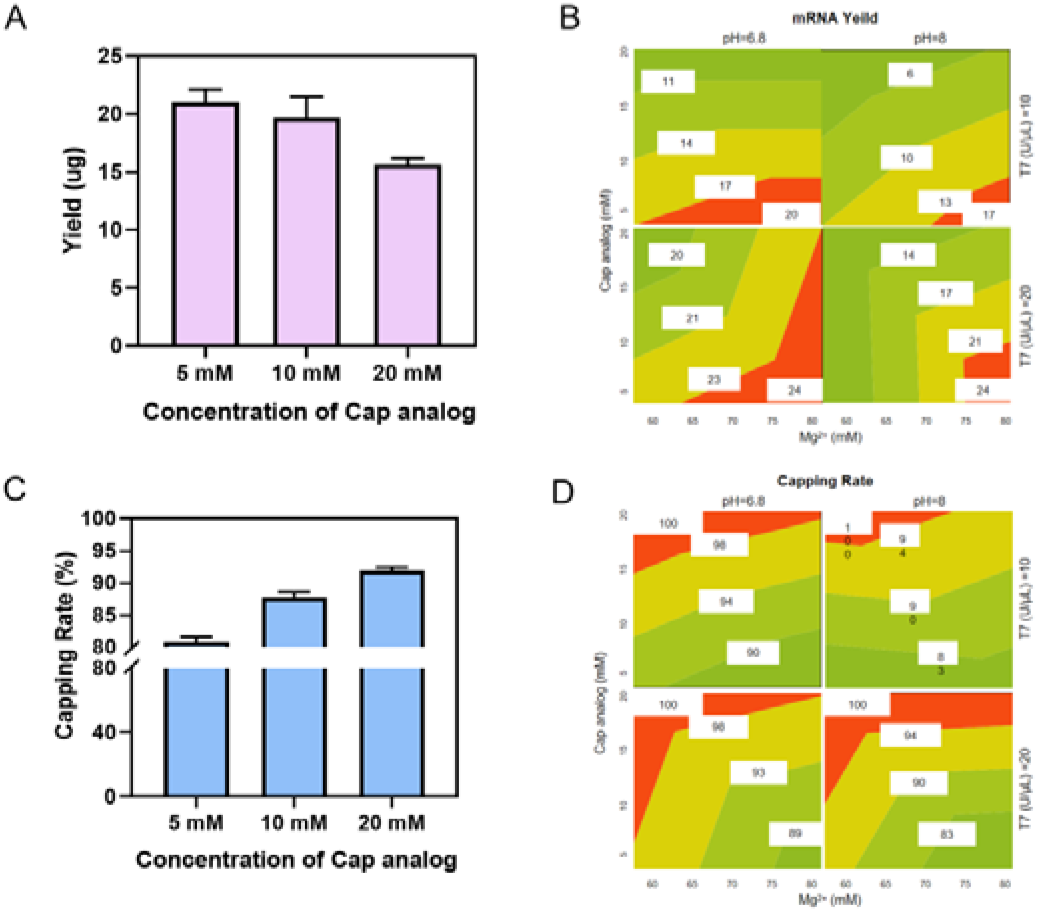
Optimizing high-yield co-transcriptional capped IVT. (a-b) Negative and positive impact of the cap analog on mRNA yield (A) and capping rate (B). A low concentration of the cap analog (5 mM) is beneficial for mRNA yield, whereas a high capping rate requires a high concentration of the cap analog (20 mM). (C-D) Response surfaces for mRNA yield (C) and capping rate (D) response surface of the DoE interaction model.

To resolve this trade-off, we proposed two mechanistically grounded strategies leveraging distinct T7 RNAP mutants. The first strategy involved engineering an Mg^2+^-insensitive T7 RNAP mutant capable of efficient transcription at low Mg:NTP ratios, thereby alleviating Mg^2+^-mediated suppression of the capping rate. The second strategy utilized a T7 RNAP mutant with enhanced affinity for cap analogs that would enable a substantial reduction in the GAG cap concentration, thereby decreasing competitive Mg^2+^ chelation, increasing the bioavailable Mg^2+^ concentration, and ultimately improving mRNA yield.

### Application of Mg^2^+-insensitive T7 RNAP for highly productive co-IVT

We established an IVT reaction under conditions of 10 mM NTP and 10 mM Mg^2+^ to screen for Mg^2+^-insensitive T7 RNAP mutants. Under these conditions, the wild-type (WT) T7 RNAP exhibited severely impaired catalytic activity, resulting in an mRNA yield of only 1 g/L. In contrast, among the 50 variants evaluated in the primary screen under identical reaction conditions, 18 displayed substantially enhanced catalytic performance relative to that of the WT, with V634T notably achieving an mRNA yield exceeding 5 g/L (Fig 2A, Table S2). Leveraging V634T as a structural and functional template, we performed a second iteration of the AI-assisted design and identified over 10 double mutants that achieved yields exceeding 5 g/L under identical conditions. The V634T/P508K double mutant emerged as the best performer and was thus selected for subsequent optimization (Fig 2B, Table S3). After three iterative rounds of screening and engineering, we obtained multiple mutants capable of sustaining maximal production, reaching an mRNA yield plateau exceeding 10 g/L (Fig 2C, Table S4). Remarkably, one of these mutants, namely V634T/P508K/C347V (hereafter designated LM9), maintained robust catalytic efficiency, achieving mRNA production yields greater than 5 g/L, even when the Mg^2+^ concentration was reduced to 5 mM (Fig 2D, Table S5). We further demonstrated that the LM9 mutant retains the critical quality attributes of therapeutic mRNA, including integrity and capping rate, while significantly reducing dsRNA content by approximately 3-fold relative to that of the WT, thereby establishing LM9 as a promising biocatalyst for therapeutic mRNA production (Fig 2E, Table S6).

**Figure 2:**
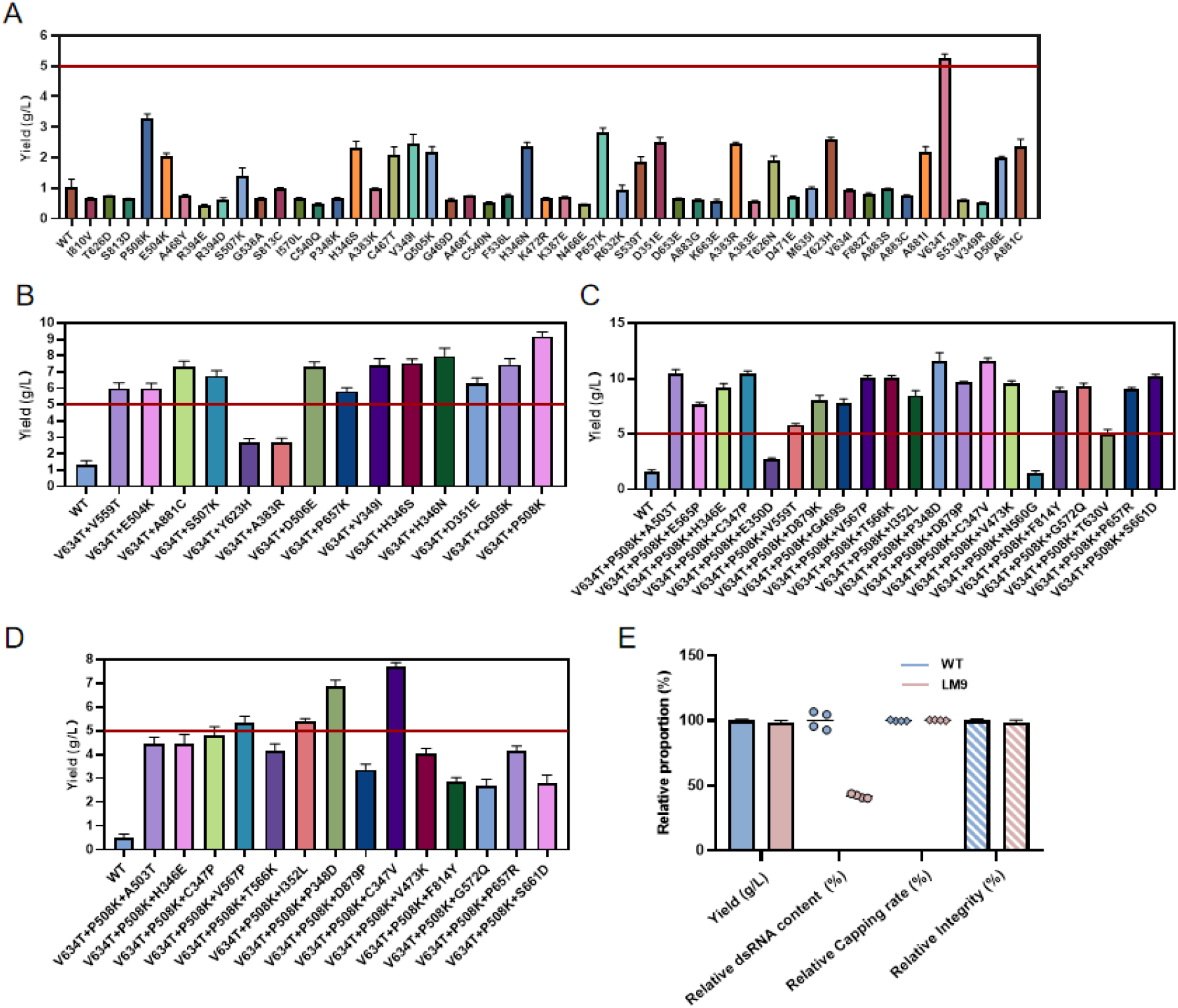
Development of Mg^2+^ -insensitive T7 RNAP. (A-C) Co-IVT yield of mRNA synthesized by the first (A), second (B) and third (C) iterations of engineered T7 RNAPs under 10 mM Mg^2+^ . (D) Co-IVT yield of mRNA synthesized by the third iteration of engineered T7 RNAPs under 5 mM Mg^2+^ . (E) Co-IVT comparison of the yield, relative dsRNA content, and capping rate of mRNA synthesized by the T7 RNAP WT and LM9 (Process 1, Table S1).

Subsequently, we used LM9 to develop a highly productive mRNA production process. We screened the Mg^2+^ requirements under the condition of 20 mM NTP. LM9 demonstrated catalytic activity at 20 mM Mg^2+^, and the mRNA yield reached 25 g/L at 40 mM Mg^2+^ (Fig 3A, Table S7). In contrast, the WT produced negligible amounts of mRNA (Fig 3B). To evaluate whether the mRNA generated by this LM9-catalyzed, highly productive co-IVT process complies with pharmaceutical quality standards, we assessed the mRNA integrity, capping rate, and dsRNA content. The mRNA product met expectations, with its integrity and capping rate demonstrating equivalence to mRNA manufactured via the canonical IVT process (Fig 3C). We found that dsRNA formation was reduced by 30-fold, which significantly exceeded the capability of LM9 (Fig 3C, Table S6).

**Figure 3:**
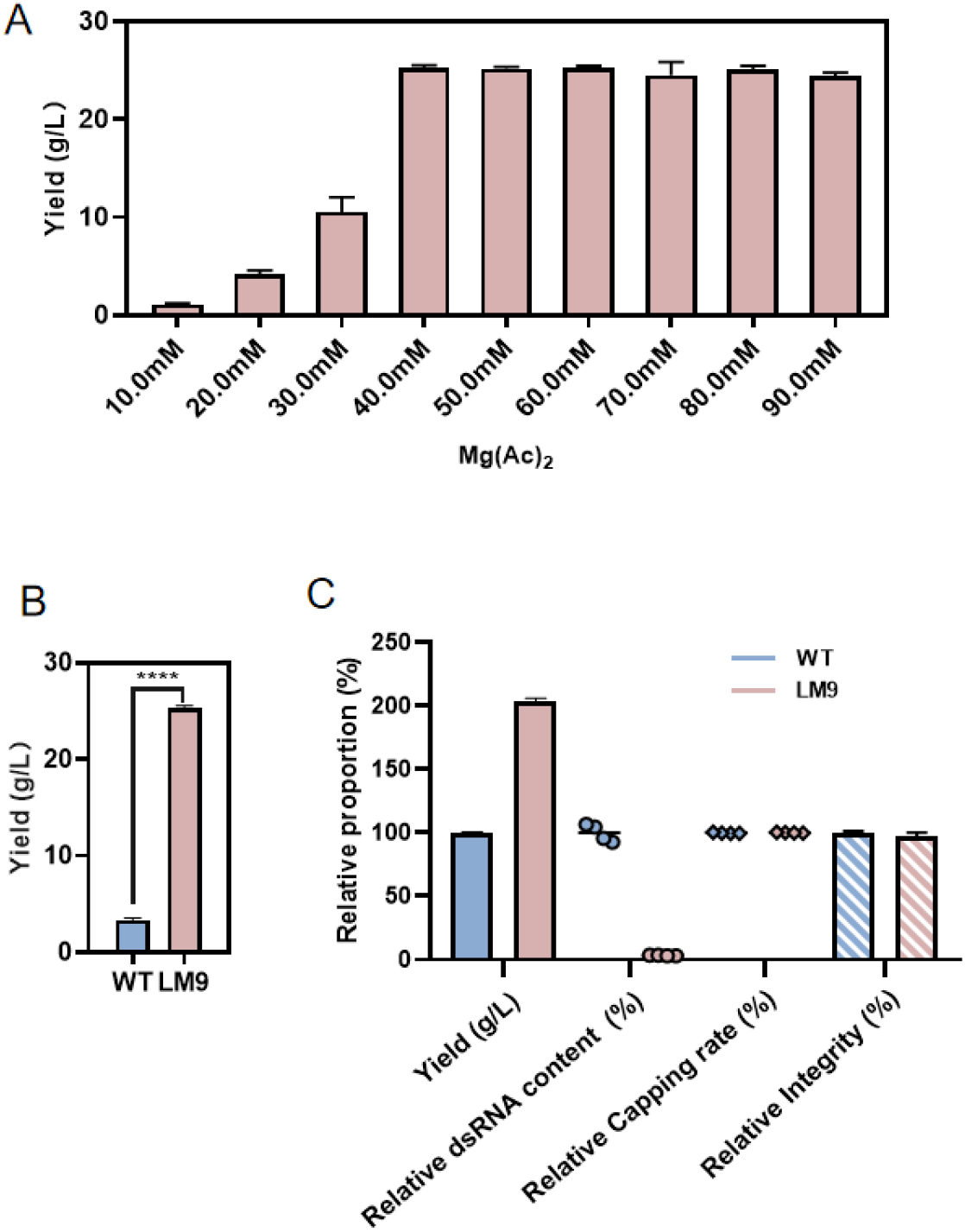
Development of a highly productive co-IVT process utilizing LM9. (A) Co-IVT yield of mRNA synthesized by LM9 across a Mg^2+^ concentration gradient. (B) Co-IVT comparison of the yield of mRNA synthesized by the T7 RNAP WT and LM9 (Process 2, Table S1). (C) Co-IVT of the yield, relative dsRNA content, and capping rate of mRNA synthesized by the T7 RNAP WT (Process 1, Table S1) and LM9 (Process 2, Table S1).

A recent study established a mathematical model for dsRNA content and initially identified NTP concentration and the Mg:NTP ratio, other than the Mg^2+^ concentration alone, as statistically significant determinants^[25]^ . We experimentally validated the significance of NTP concentration and extended its applicability to NTP concentrations up to 20 mM. At a fixed Mg:NTP molar ratio of 4:1, increasing NTP concentrations positively correlated with IVT yield and negatively with dsRNA content (Fig S3, Table S8).

### Application of T7 CapMax RNAP for highly productive co-IVT

The commercially available T7 CapMax RNAP exhibits superior capping efficiency. We initially evaluated the performance of CapMax across a GAG cap concentration gradient. Relative to the WT, which required a GAG:GTP ratio of 0.8 to achieve a capping rate exceeding 99%, CapMax achieved an equivalent capping rate at a GAG:GTP ratio of 0.06 (Fig 4A, B, Table S9), reducing cap analog consumption by over 90%. The T7 CapMax RNAP was then employed to develop an alternative, highly productive co-IVT process.

**Figure 4:**
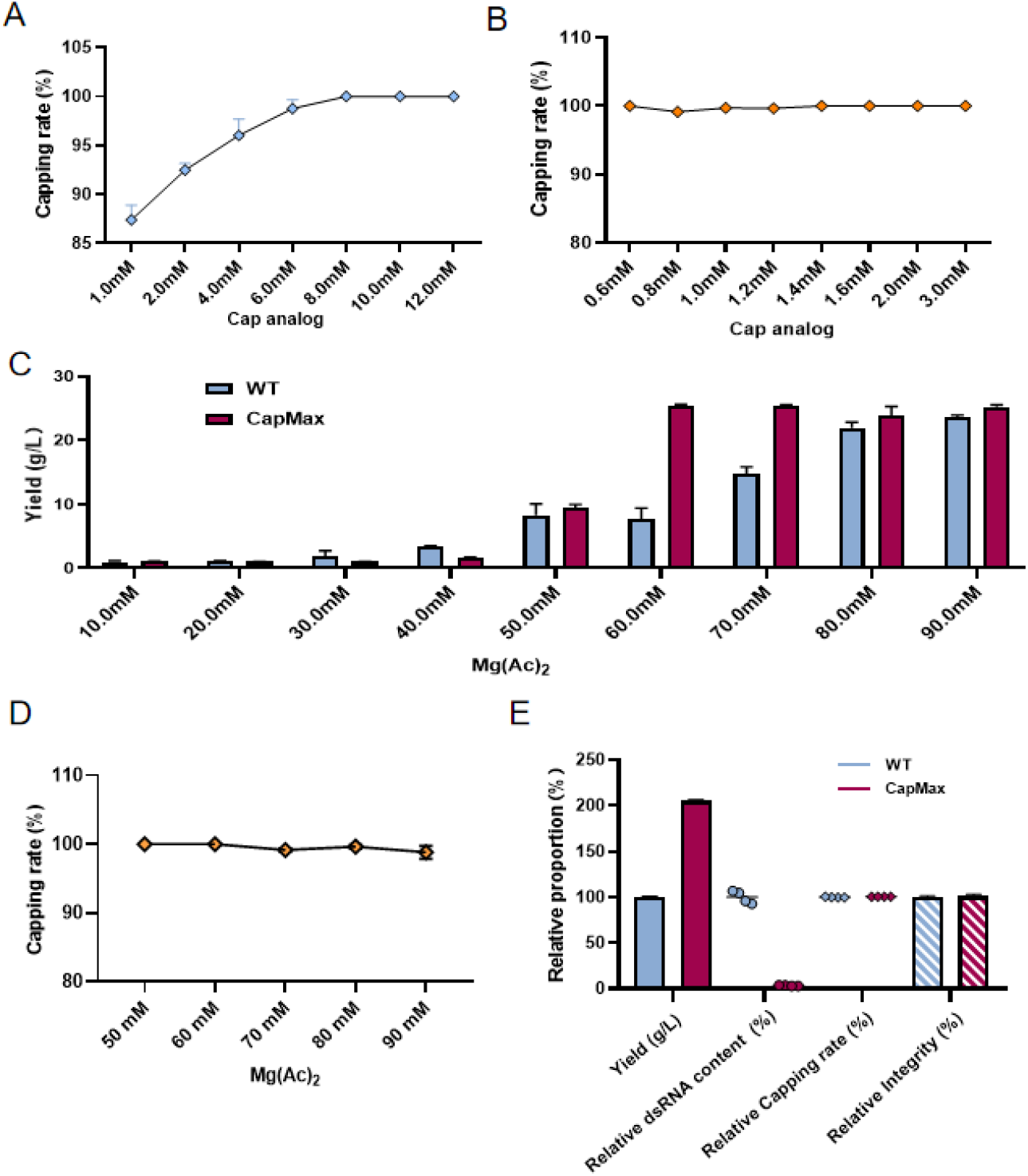
Development of a highly productive co-IVT utilizing CapMax. (A-B) Comparative analysis of capping rates between the WT (A) and CapMax (B) across a concentration gradient of GAG cap. (C) Co-IVT yield of mRNA synthesized by the WT and CapMax across a concentration gradient of Mg^2+^. (D) Analysis of capping rates catalyzed by CapMax across a concentration gradient of GAG cap. (E) Comparison of the Co-IVT yield, dsRNA content and capping rate of mRNA catalyzed CapMax (Process 3, Table S1) compared with those of the WT (Process 1, Table S1).

The Mg^2+^ requirement was systematically evaluated across a concentration range (10–90 mM), with the NTPs and GAG cap concentrations fixed at 20 mM. The results demonstrated that CapMax exhibits unexpectedly reduced Mg^2+^ dependence under the highly productive co-IVT conditions, achieving a maximal mRNA yield at 60 mM Mg^2+^, in contrast to the WT, which required 80 mM Mg^2+^ to sustain comparable performance (Fig 4C, Table S7). Moreover, a high capping rate exceeding 99% was maintained across the tested Mg^2+^ concentration range (Fig 4D). To assess whether CapMax exhibits broad Mg^2+^-insensitivity, the aforementioned Mg^2+^ requirement evaluation was performed in absence of the cap analog and using the DNA template with promoter +1′ substituted with G, which is suitable for non-co-IVT. Under these conditions, CapMax exhibited a stronger Mg^2+^ utilization efficiency relative to that of the WT, with an optimal Mg^2+^ concentration range narrowed to 60–80 mM, compared with the 50–90 mM range exhibited by the WT (Fig S4, Table S10). These results indicate that the reduced Mg^2+^ requirement of CapMax under highly productive co-IVT conditions arose indirectly from its significantly enhanced affinity for cap analogs. CapMax did not exhibit comprehensive Mg^2+^ insensitivity.

Subsequent Mg^2+^ titration further confirmed that 54 mM is sufficient for CapMax to catalyze highly productive co-IVT, reaching a yield plateau (Fig S5, Table S7). Subsequent GAG cap titration experiments demonstrated that a capping rate exceeding 99% was maintained even when the GAG cap concentration was reduced to 2 mM (corresponding to a GAG:GTP ratio of 0.1), whereas the mRNA yield decreased when the GAG cap concentration fell below 1 mM (Fig S6, Table S11). A comparison with canonical co-IVT was then performed, and the results revealed comparable mRNA integrity and capping rates, along with a significant reduction in dsRNA content (Fig 4E, Table S6).

Two distinct and highly productive co-IVT processes have been successfully established. To determine which process has greater potential for scalable development, a comparative analysis was performed under distinct optimized conditions. Both processes achieved comparably high yields, along with comparable mRNA integrity, capping rates, and dsRNA content (Table 1). However, the GAG cap consumption significantly differed between the two processes. Although the optimal GAG cap concentration for LM9 has yet to be determined, CapMax exhibited a markedly higher utilization of GAG cap, thereby conferring a distinct advantage in the CoGs.

**Table 1:** Comparative analysis of the two highly productive co-IVT processes.

| RNAP | LM9 | CapMax |
| --- | --- | --- |
| Total NTP | 80 mM | 80 mM |
| GAG cap | 20 mM | 1 mM |
| Magnesium acetate | 40 mM | 54 mM |
| Yield | 25.32 ± 0.24 g/L | 25.06 ± 0.62 g/L |
| Capping rate | 99.66 ± 0.30% | 100% |
| dsRNA content | 66.87±5.14 ppm | 64.74±7.75 ppm |
| mRNA integrity | 83.78±2.35 % | 86.60±1.95% |

### Optimization of the highly productive co-IVT process

Sequence-adapted NTP supplementation maximizes the NTP conversion rate and thereby enhances IVT productivity^[27]^. We validated this strategy and achieved an mRNA yield reaching 26.07±0.42 g/L (Fig 5A). This solution, along with all the aforementioned highly productive co-IVT mixtures, exhibited a gel-like rheological behavior, rendering it nearly non-pourable (Fig S7). In laboratory-scale experiments, dilution with a 10-fold volume of water followed by incubation at 37°C for 10 min fully solubilized the gel-like IVT mixture, enabling subsequent purification and analytical characterization. However, this dilution-based approach is impractical for large-scale manufacturing due to insufficient space volume in the bioreactor. Consequently, alternative solubilization strategies must be developed.

**Figure 5:**
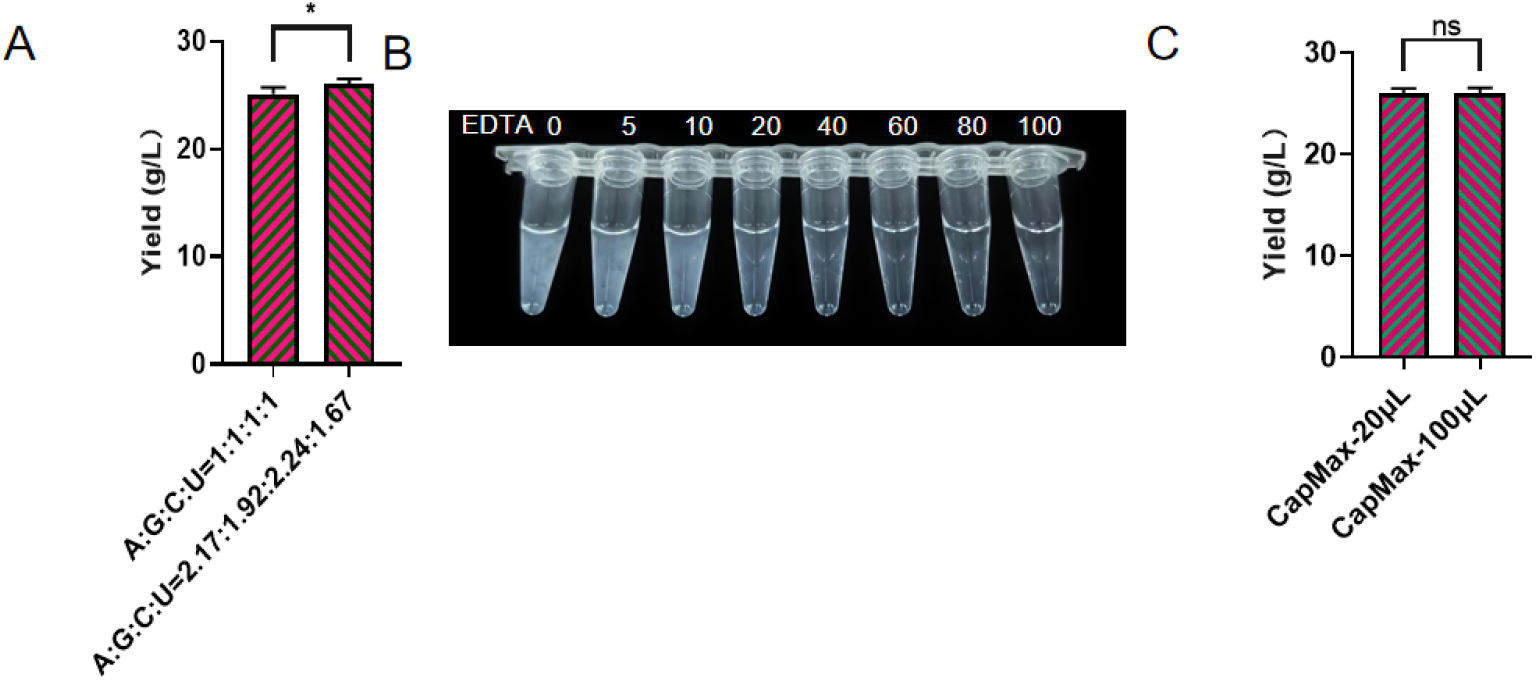
Optimization of a highly productive co-IVT process. (A) Sequence-adapted NTP supplementation enhanced mRNA yield (Process 11, Table S1). (B) Visualization of the appearance and solubility status of the highly productive co-IVT product with or without EDTA treatment (Process 12, Table S1). (C) Co-IVT yield following EDTA treatment.

As the gel-like material likely arises from magnesium-mediated RNA polymerization, resulting in a viscoelastic nucleic acid–metal hydrogel^[28]^, introducing EDTA to chelate Mg^2+^ in the IVT mixture represents a highly effective strategy for solubilizing the RNA hydrogel and restoring the IVT mixture to a free-flowing liquid state^[28]^ . We determined that a final disodium EDTA concentration of 60 mM fully resolved the hydrogel and restored the IVT reaction mixture to a free-flowing state (Fig 5B). Subsequent analytical assessments confirmed that critical quality attributes, including yield integrity, capping rate, and dsRNA content, were unaffected by EDTA treatment (Fig 5C).

### Scale-up verification

To showcase the industrial utility of the highly productive co-IVT process, we manufactured mRNA at a 10-mL reaction scale, followed by DNase I digestion, oligo(dT)-based affinity chromatography, and tangential flow filtration, in accordance with the most widely applied mRNA drug substance manufacturing protocols. We confirmed that the EDTA-mediated solubilization strategy remains effective at the 10-mL scale (Table 2). Furthermore, characterization of EDTA-treated samples demonstrated that the yield, integrity, capping rate, and dsRNA content were statistically equivalent between the 100-μL and 10-mL scales, indicating the absence of scale-dependent effects (Table 2). These critical quality and key performance attributes aligned with predefined targets, confirming that the significantly enhanced co-IVT productivity did not compromise downstream purification performance. The present, highly productive co-IVT process demonstrates robust compatibility and plug-and-play integration, enabling seamless incorporation into established mRNA manufacturing workflows (Fig S9).

**Table 2:** Comparative analysis of the yield, integrity, capping rate, and dsRNA content of the highly productive co-IVT product at different scale.

| Substitutions | Scale | Yield of mRNA products (g/L) | Relative dsRNA content (%) | Capping rate (%) | integrity (%) |
| --- | --- | --- | --- | --- | --- |
| T7 CapMax-60mM EDTA | 100 μL | 26.02±0.50 | 62.09±4.58 | 100.00±0.00 | 85.15±1.54 |
| T7 CapMax-60mM EDTA | 10 mL | 25.85±0.67 | 65.78±3.33 | 100.00±0.00 | 84.53±0.75 |

## DISCUSSION

Herein, we report a novel, highly productive co-IVT process that achieves an unprecedented mRNA yield approximately 3-fold higher than that of the current industry average, without compromising critical quality attributes. The process maintains an identical input of T7 RNAP and DNA template yet significantly reduces cap analog consumption. As these three reagents constitute over 80% of the total raw material cost of co-IVT, the present process markedly improves the economic viability of mRNA manufacturing.

The optimized, highly productive co-IVT process achieved an mRNA yield of 26.13 g/L, which is equivalent to an almost complete conversion of 80 mM NTPs into mRNA product. The observed yield ceiling was imposed by the maximum concentrations of commercially available NTP stock solutions, not by the intrinsic capability of the process or by substrate or product solubility limitations. By enabling total NTP concentrations of 100–120 mM in the reaction mixture using lyophilized NTPs, we anticipate that an mRNA yield exceeding 35 g/L could be achieved.

The most striking characteristic of the reaction under highly productive conditions was the transition of the product solution from a free-flowing liquid to a hydrogel. EDTA treatment efficiently solubilized the hydrogel, thereby enabling product recovery from the reactor during large-scale manufacturing. The process was successfully validated at a 10-mL scale, demonstrating its scalability and compatibility with downstream purification procedures. However, further process scale-up presents a potential risk of failure, as excessive solution viscosity may impair mixing efficiency in wave bioreactors, resulting in substantial mass-transfer limitations and a consequent decline in overall productivity.

Moreover, we introduce two distinct engineered T7 RNAPs. LM9 retains high catalytic activity under conditions of an extremely low Mg:NTP molar ratio, at which the WT enzyme is inactive and free Mg^2+^ is undetectable^[29]^. The significance of free Mg^2+^ likely stems from its role in binding to the Mg^(A)^ site and stabilizing productive substrate binding at the P site of T7 RNAP. Although T7 RNAP catalyzes the de novo RNA synthesis of MgNTP^2−^ via a two-metal mechanism, the Mg^2+^ delivered by the initiating nucleotide occupies a distinct pocket formed by the β-and γ-phosphates of the initiating nucleotide and surrounding basic residues of T7 RNAP, distinct from the canonical catalytic Mg^(A)^ site^[30]^. Therefore, we sought to elucidate the mechanistic basis underlying the ability of Mg^2+^-insensitive mutants, including LM9, to efficiently catalyze IVT in the absence of free Mg^2+^. The mutated residues are spatially dispersed throughout the active pocket, rather than clustered within a localized subregion (Fig S). Based on these structural insights, we propose that magnesium-insensitive mutants restore transcriptional activity by modulating the conformational dynamics of the active-site pocket, thereby facilitating the productive binding of phosphate-chelated Mg^2+^ at Site A and enabling the nucleophilic attack essential for transcription initiation (Fig S10). Mg^2+^ concentration represents the most critical process parameter for every critical quality attribute of IVT and co-IVT. However, we found that a substantial reduction in Mg^2+^ concentration failed to improve either the capping rate or transcript integrity, yielding only a marginal decrease in dsRNA levels, contrary to extensive prior knowledge. Meanwhile, simultaneously increasing Mg^2+^ and NTP concentrations did not impair IVT product quality attributes. Our findings indicate that the free Mg^2+^ concentration, rather than the total Mg^2+^ concentration, is the critical process parameter for IVT.

CapMax, a rationally engineered T7 RNAP with enhanced affinity for cap analogs, recapitulates the functional properties of LM9, exhibiting notably robust co-IVT efficiency under low Mg:NTP ratios. Critically, this feature is abolished in the absence of a cap analog, and the ratio threshold for activity is substantially higher than that for LM9, indicating a mechanistically distinct mode of action. Assembly of the productive transcription initiation complex comprising T7 RNAP, template DNA, initiating NTP substrates, and Mg^2+^ represents the rate-limiting step in IVT reactions. We propose that CapMax promotes co-IVT initiation complex formation by enhancing affinity for cap analogs, thereby reducing the dependence on free Mg^2+^.

Both LM9 and CapMax display previously uncharacterized functional properties that expand the operational parameter space for IVT and co-IVT. Beyond the highly productive system reported herein, these mutants present opportunities for broader RNA synthesis applications.

## AUTHOR CONTRIBUTIONS

Wei He: Methodology, Validation, Formal Analysis, Visualization, Writing – Original Draft. Kunkun Zhang: Methodology, Formal Analysis, Visualization, Writing – Original Draft. Guiying Ji: Investigation. Hongli Zhou: Investigation. Ya Dai: Investigation. Minghao Xu: Investigation, Resources. Xiaoyu Xu: Supervision, Funding Acquisition. Qiuheng Jin: Conceptualization, Writing – Review & Editing, Supervision, Project Administration.

## REFERENCES

1. Xu B, Sun Z and Rokita SE. A split gene approach to alleviate severe inhibition of catalysis by substrate. Biochemistry. 2025; 64: 2867–2876.

2. Ma Y, Liu N, Greisen P et al. Removal of lycopene substrate inhibition enables high carotenoid productivity in yarrowia lipolytica. Nat Commun. 2022; 13: 572.

3. Zhao F, Zhou Z, Dang Y et al. Genome-wide role of codon usage on transcription and identification of potential regulators. Proc Natl Acad Sci USA. 2021; 118: e2022590118.

4. Völler JS. Locking in lipases. Nat Catal. 2025; 8: 507.

5. Rudroff F, Mihovilovic MD, Gröger H et al. Opportunities and challenges for combining chemo- and biocatalysis. Nat Catal. 2018; 1: 12–22.

6. Ulusu NN. Evolution of enzyme kinetic mechanisms. J Mol Evol. 2015; 80: 251–257.

7. Prywes N, Phillips NR, Tuck OT et al. Rubisco function, evolution, and engineering. Annu Rev Biochem. 2023; 92: 385–410.

8. Zhang Y, Zhang G, Wang T et al. Understanding cytochrome p450 enzyme substrate inhibition and prospects for elimination strategies. ChemBioChem. 2024; 25: e202400297.

9. Smith BT, Knutsen JS and Davis RH. Empirical evaluation of inhibitory product, substrate, and enzyme effects during the enzymatic saccharification of lignocellulosic biomass. Appl Biochem. Biotechnol. 2010; 161: 468–482.

10. Jha RK, Narayanan N, Pandey N et al. Sensor-enabled alleviation of product inhibition in chorismate pyruvate-lyase. ACS Synth Biol. 2019; 8: 775–786.

11. Guo L, Liu Z, Song S et al. Maximizing the mRNA productivity for in vitro transcription by optimization of fed-batch strategy. Biochem Eng J. 2024; 210: 11.

12. Zak MM and Zangi L. Clinical development of therapeutic mRNA applications. Mol Ther. 2025; 33: 2583–2609.

13. Weber JS, Carlino MS, Khattak A et al. Individualised neoantigen therapy mRNA-4157 (v940) plus pembrolizumab versus pembrolizumab monotherapy in resected melanoma (keynote-942): A randomised, phase 2b study. Lancet. 2024; 403: 632–644.

14. Sahin U, Schmidt M, Derhovanessian E et al. Individualized mRNA vaccines evoke durable t -cell immunity in adjuvant tnbc. Nature. 2026; 651: 1088–1096.

15. Intellia heads to fda with first in vivo crispr-based gene editing therapy. Nat Biotechnol. 2026; 44: 676.

16. Koeberl D, Schulze A, Sondheimer N et al. Interim analyses of a first-in-human phase 1/2 mRNA trial for propionic acidaemia. Nature. 2024; 628: 872–877.

17. Moschioni M, Siraji RA, Dissard R et al. mRNA vaccines and therapeutics beyond COVID-19: A review of the global clinical development landscape, low- and middle-income countries involvement and relevance to their contexts. Hum Vaccin Immunother. 2026; 22: 2628424.

18. Yisraeli JK and Melton DA. Synthesis of long, capped transcripts in vitro by SP6 and T7 RNA polymerases. Methods Enzymol. 1989; 180: 42–50.

19. Ghosh A and Lima CD. Enzymology of RNA cap synthesis. WIREs RNA. 2010; 1: 152–172.

20. Henderson JM, Ujita A, Hill E et al. Correction: Cap 1 messenger RNA synthesis with co-transcriptional cleancap® analog by in vitro transcription. Curr Protoc. 2021; 1: e336.

21. Jain A and Vale RD. RNA phase transitions in repeat expansion disorders. Nature. 2017; 546: 243–247.

22. Kis Z, Kontoravdi C, Shattock R et al. Resources, production scales and time required for producing RNA vaccines for the global pandemic demand. Vaccines (Basel). 2020; 9: 3.

23. Skok J, Meguar P, Vodopivec T et al. Gram-scale mRNA production using a 250-ml single-use bioreactor. Chem Ing Tech. 2022; 94: 1928–1935.

24. Stover NM, Ahmadi S, Rosenfeld J et al. Model-based optimization of fed-batch in vitro transcription. ChemBioChem. 2025; 26: e202500485.

25. Boman J, Marusic T, Seravalli TV et al. Quality by design approach to improve quality and decrease cost of in vitro transcription of mRNA using design of experiments. Biotechnol Bioeng. 2024; 121: 3415–3427.

26. Team B. Geoflow-v2: A unified atomic diffusion model for protein structure prediction and de novo design. bioRxiv, 10.1101/2025.05.06.652551, May11, 2025, pre-print: not peer-reviewed.

27. Zhao K, Raffaele J, Holland DA et al. Enhancing mRNA vaccine production: Optimization of in vitro transcription for improved yield and purity. Biotechnol Prog. 2026; 42: e70109.

28. Yewdall NA, André AAM, van Haren MHI et al. Atp:Mg2+ shapes material properties of protein–RNA condensates and their partitioning of clients. Biophys J. 2022; 121: 3962–3974.

29. Marusic T, Faganel T and Sekirnik R. High-throughput monitoring of free mg2+ in ivt reaction reveals effects of free mg^2+^ on mRNA synthesis and DNA digestion. Anal Bioanal Chem. 2026; 418: 3983–3997.

30. Yin YW and Steitz TA. The structural mechanism of translocation and helicase activity in T7 RNA polymerase. Cell. 2004; 116: 393–404.

