## Supplementary Figures for "Maximizing the productivity of co-transcriptional in vitro transcription"

Figure Captions

**Figure S1.** Schematic of the DNA template.

**Figure S2.** Effect summary of DoE assay on mRNA yield, capping rate and dsRNA.

**Figure S3.** (A) mRNA yield and (B) dsRNA content in co-IVT reactions catalyzed by T7 RNAP WT, as a function of magnesium cofactor concentration.

**Figure S4.** mRNA yield in IVT reactions catalyzed by T7 RNAP WT and Capmax, as a function of magnesium cofactor concentration.

**Figure S5.** mRNA yield in co-IVT reactions catalyzed by T7 RNAP WT and Capmax, as a function of magnesium cofactor concentration.

**Figure S6.** mRNA yield in co-IVT reactions catalyzed by T7 RNAP WT and Capmax, as a function of cap analog concentration.

**Figure S7.** Visualization of the appearance and solubility status of the highly productive co-IVT product without EDTA treatment.

**Figure S8.** Comparison of mRNA yield, dsRNA content, capping efficiency, and integrity of co-IVT products catalyzed by T7 RNAP CapMax at 100 μL and 10 mL reaction scales.

**Figure S9.** Flowchart of mRNA in vitro production and purification at scaled-up reaction condition.

**Figure S10.** Summary of T7 RNAP mutation sites that confer insensitivity to the cofactor magnesium during in vitro transcription.

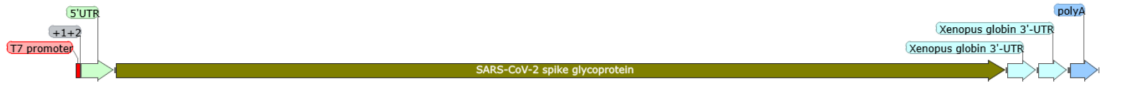

Figure S1. Schematic of the DNA template.

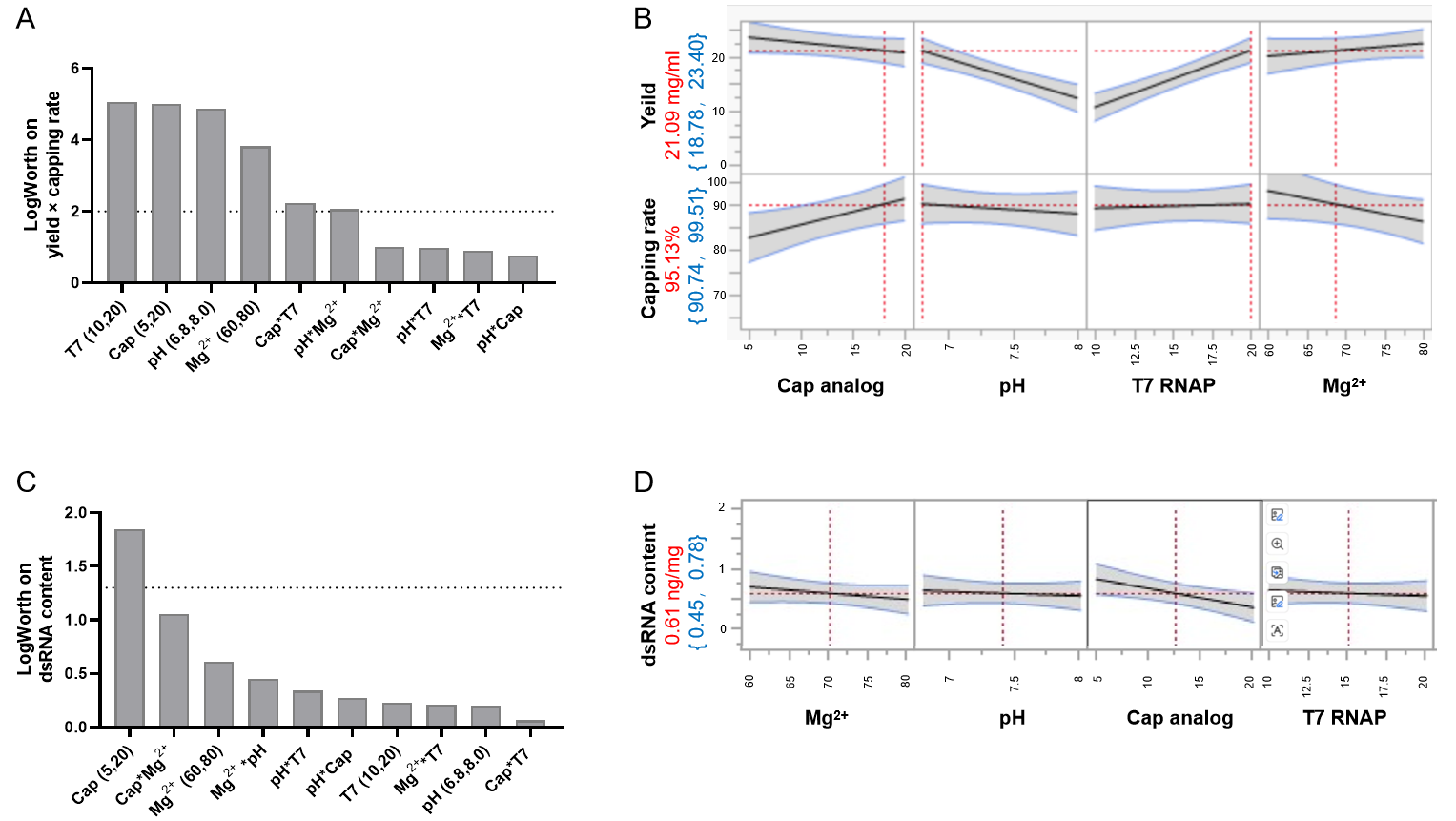

Figure S2. Effect summary of DoE assay on mRNA yield, capping rate and dsRNA. (A) Effect summary on mRNA yield combining capping rate, which showed T7 RNAP, cap analog, pH and Mg2+ are the key factor. (B) Prediction profiler of mRNA yield and capping rate. Under the optimal condition, the mRNA yield could reach to 21.09 mg/ml, meanwhile the capping rate can be 95.13. (C) Effect summary on byproduct dsRNA content. (D) Prediction profiler of dsRNA content.

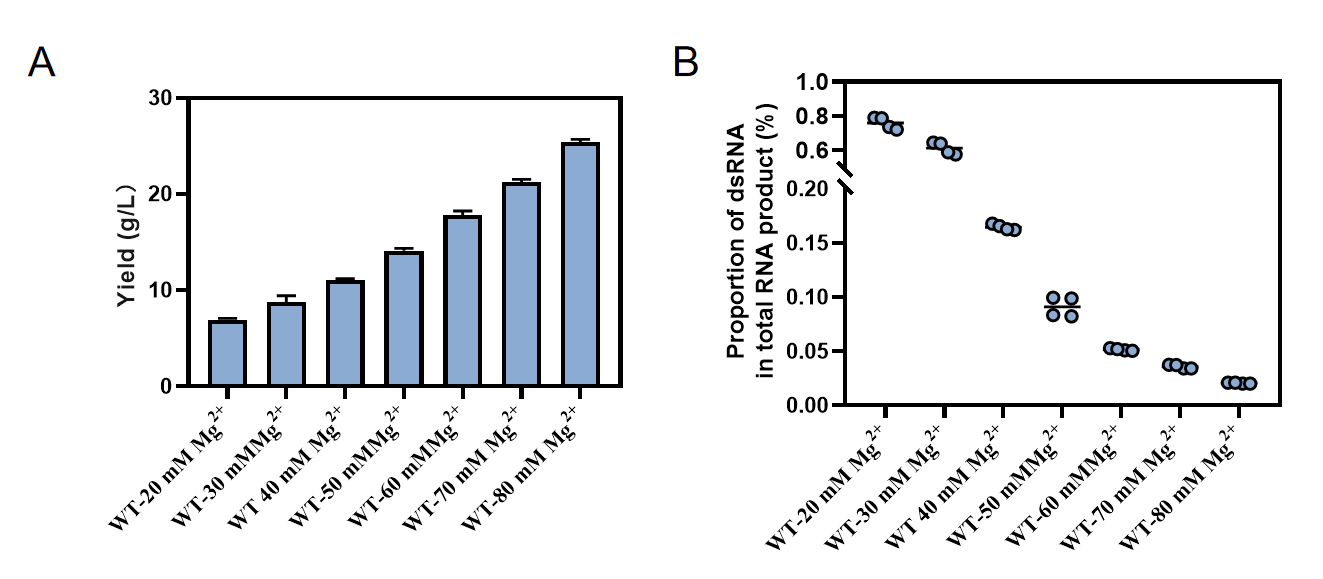

Figure S3. (A) mRNA yield and (B) dsRNA content in co-IVT reactions catalyzed by T7 RNAP WT, as a function of magnesium cofactor concentration

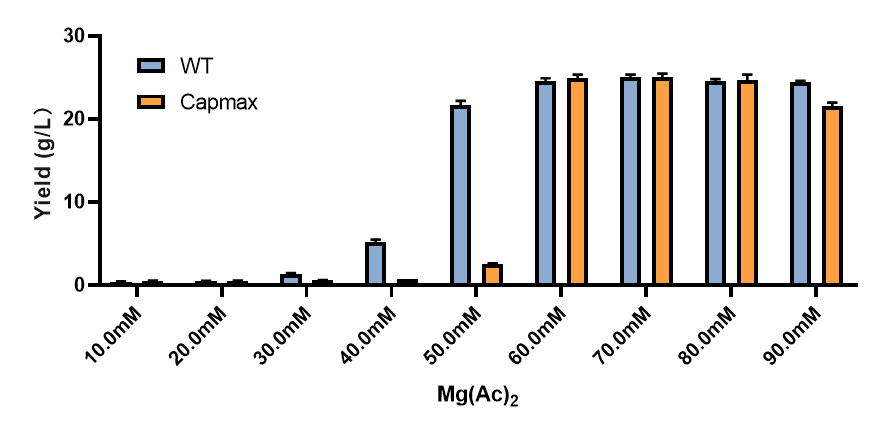

Figure S4. mRNA yield in IVT reactions catalyzed by T7 RNAP WT and Capmax, as a function of magnesium cofactor concentration..

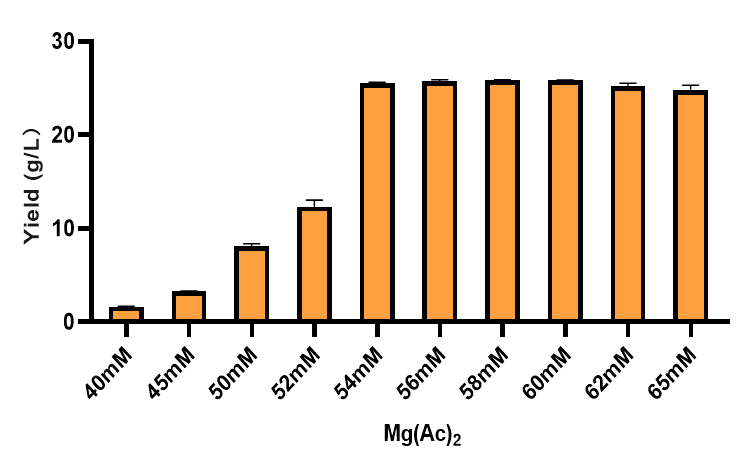

Figure S5. mRNA yield in co-IVT reactions catalyzed by T7 RNAP WT and Capmax, as a function of magnesium cofactor concentration.

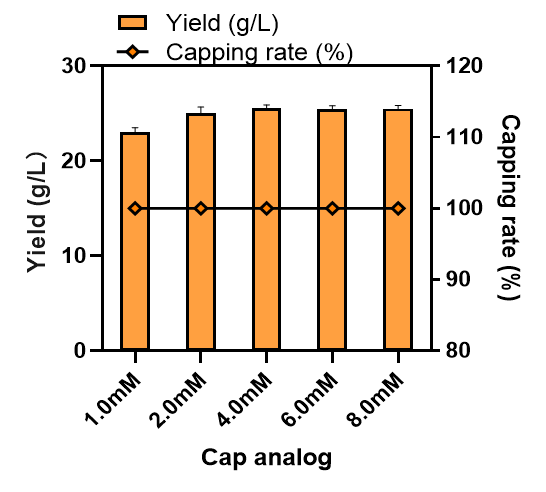

Figure S6. mRNA yield in co-IVT reactions catalyzed by T7 RNAP WT and Capmax, as a function of cap analog concentration

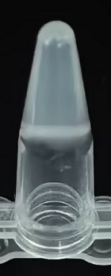

Figure S7. Visualization of the appearance and solubility status of the highly productive co-IVT product without EDTA treatment..

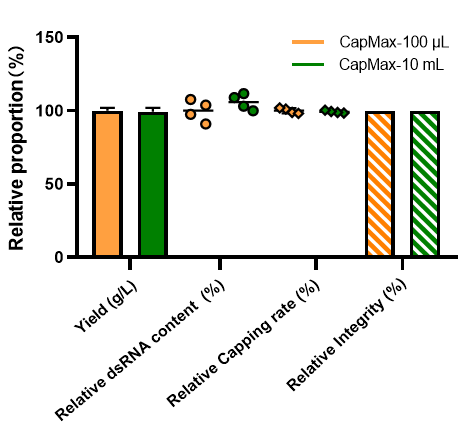

Figure S8. Comparison of mRNA yield, dsRNA content, capping efficiency, and integrity of co-IVT products catalyzed by T7 RNAP CapMax at 100 μL and 10 mL reaction scales.

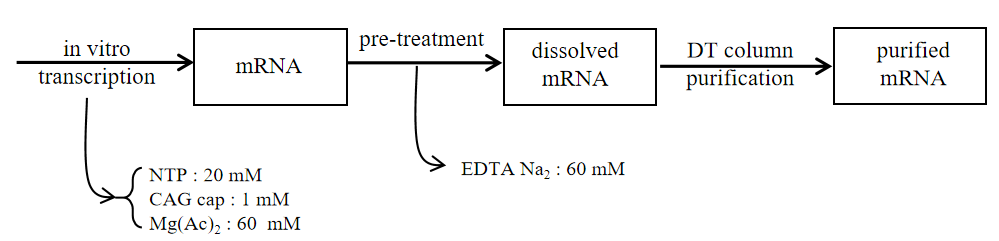

Figure S9. Flowchart of mRNA in vitro production and purification at scaled-up reaction condition

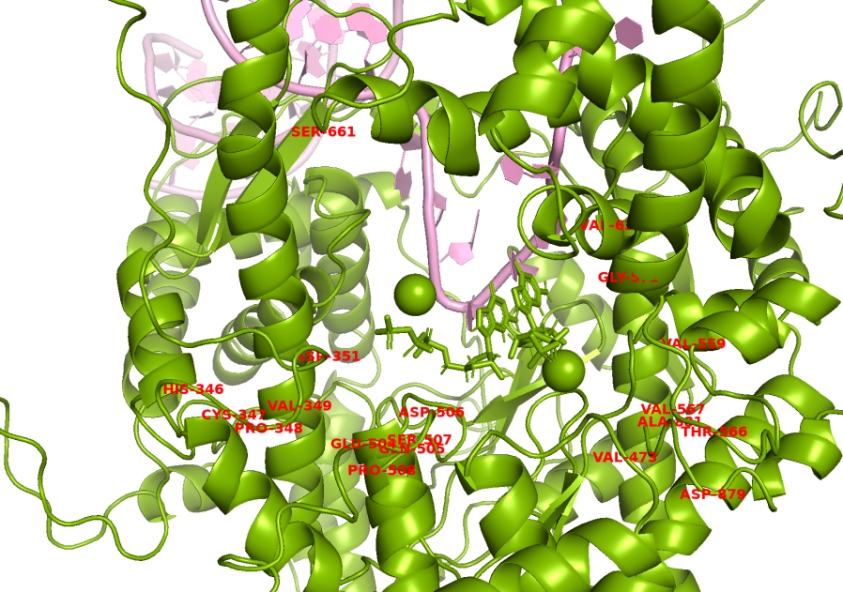

Figure S10. Summary of T7 RNAP mutation sites that confer insensitivity to the cofactor magnesium during in vitro transcription.
